# Streamlining Spatial Transcriptomics for Human Kidney Tissue

**DOI:** 10.1101/2025.09.02.673834

**Authors:** Son Vo, Kieran Meadows, Han Do, Kevin Chapman, Graciela Andonegui, Daniel A. Muruve, Thang Pham, Justin Chun

## Abstract

Single-cell spatial technologies have emerged in recent years, enabling tissue architecture and organization characterization at unprecedented resolution. However, computational analysis of spatial transcriptomics data is often a bottleneck for scientific discoveries in the absence of a dedicated bioinformatician. Here, we describes a workflow to annotate cell types from a dataset generated using NanoString’s CosMx single-cell resolution spatial transcriptomic technology, enabling a comparison between healthy kidney biopsies and diseased tissue. We validated our pipeline’s accuracy with both gene expression analysis and pathological changes associated with biopsy-proven diabetic kidney disease (DKD). Through precise cell type annotation, we observed significant changes in the proportions of podocytes and immune cells in DKD, with DKD tissue showing regional enrichment of immune cells and differential gene expression. Notably, injured proximal tubules had the expected increased expression of *HAVCR1* and *VCAM1* and genes associated with diabetes, including *IL18*, *ITGA3* and *ITGB1*. The entire workflow, now fully integrated into the BioTuring SpatialX (Lens V2.0), is available as a platform designed for users with no formal bioinformatics training.

## Introduction

The kidney is a complex organ consisting of over 30 specialized cells, including various epithelial cell types, endothelial cells, interstitial cells, and immune cells (1,2). Injury or disease of the kidneys can induce changes to the cellular microenvironment and kidney architecture, initiating inflammatory and fibrotic processes to stimulate repair and regeneration. Sustained pathological processes can lead to permanent injury, leading to chronic kidney disease (CKD), a global burden impacting approximately ten percent of the world’s population (3). Diabetic kidney disease (DKD) remains the leading cause of kidney disease worldwide, but the complex pathophysiology still remains poorly characterized (4,5).

The histopathological features of DKD, based on the Renal Pathology Society, are primarily based on glomerular lesions that are distinct from chronic changes of tubular atrophy and interstitial fibrosis commonly seen in biopsies of patients with CKD (6). However, the cell-specific gene expression changes in regions of diseased tissue specific to DKD are difficult to ascertain due to the lack of spatial resolution using conventional experimental methodologies. Recently, Abedini et al. demonstrated the power of single-cell resolution spatial transcriptomics in analyzing healthy and DKD biopsies (1). Utilizing data generated using NanoString’s CosMx spatial molecular imaging, subclustering analysis of stromal cells distinguished 12 different cell subtypes that separated different myofibroblast types and discriminated between vascular smooth muscle cells from myofibroblasts. CosMx spatial molecular imaging technology enables high-resolution, transcript-level mapping of gene expression in tissues, while preserving spatial context at single-cell resolution. This technology is particularly valuable for studying complex organs such as the kidney, where cellular diversity and spatial organization are critical to understanding both health and disease.

Accurate and standardized cell type annotation in CosMx data remains a significant challenge, especially in diseased tissues due to the complexity of cellular states and pathological changes. Therefore, we sought to introduce an improved workflow for annotating cell types of kidney tissue in CosMx data. Using two healthy kidney samples and two biopsy-proven type 2 diabetic kidney disease samples, cell types within the glomerulus and nephron tubular system were categorized into 8 clusters. Additionally, immune cell types in DKD were identified, thus enhancing our understanding of the cellular and spatial changes associated with diabetic kidney disease.

## Methods

### CosMx sample preparation for kidney biopsies

Kidney biopsy samples were obtained from the University of Calgary Precision Medicine in Nephrology Program’s Biobank for the Molecular Classification of Kidney Disease (REB17-0639). Patient samples were approved by the Conjoint Health Research Ethics Board at the University of Calgary. Human kidney biopsies from control were healthy tissue from protocol kidney biopsies pre-transplantation and patients with diabetic kidney diseases. Formalin-fixed, paraffin-embedded samples were prepared according to the manufacturer’s specifications (NanoString Technologies). In brief, tissue sections were cut at 5 µm thickness using a Leica microtome onto Leica BOND PLUS slides (Leica Biosystems, S21.2113.A). Via the NanoString CosMx spatial molecular imager (SMI) Technology Access Program, a 1000-plex CosMx Human Technology Access Program (TAP) RNA panel was performed with the addition of kidney-specific custom genes: *APOL1, AQP1, AQP2, CLDN10, EMCN, HAVCR1, LRP2, NOS1, NPHS1, PALMD, PIEZO2, PLVAP, PROM1, PROX1, REN, SCN2A, SH3GL3, SLC12A1, SLC12A3, SLC14A2, SLC26A4, SLC26A7, SLC4A9, SLC5A12, SLC5A2, SLC7A13, SLC8A1, SYN1, UNC5D*.

Morphology visualization was performed using the markers CD298/B2M, PanCK, CD45, CD68, and DAPI upfront stain and CD10 and podoplanin post stain. Slides were imaged using the CosMx SMI.

### Preprocessing and cell type annotation

The full pipeline is summarized in Figure 1. In brief, pre-processing quality control was conducted by assigning cell transcripts to cell barcodes and subsequent data normalization. Subsequent batch variability was identified using principal component analysis (PCA) and corrected using single-cell variational inference (scVI) (7). Samples were then visualized using UMAP analysis and clustered using Leiden clustering. Subsequent manual cell type annotation was performed using established hallmark cell type genes for kidney tissue.

**Figure 1.**
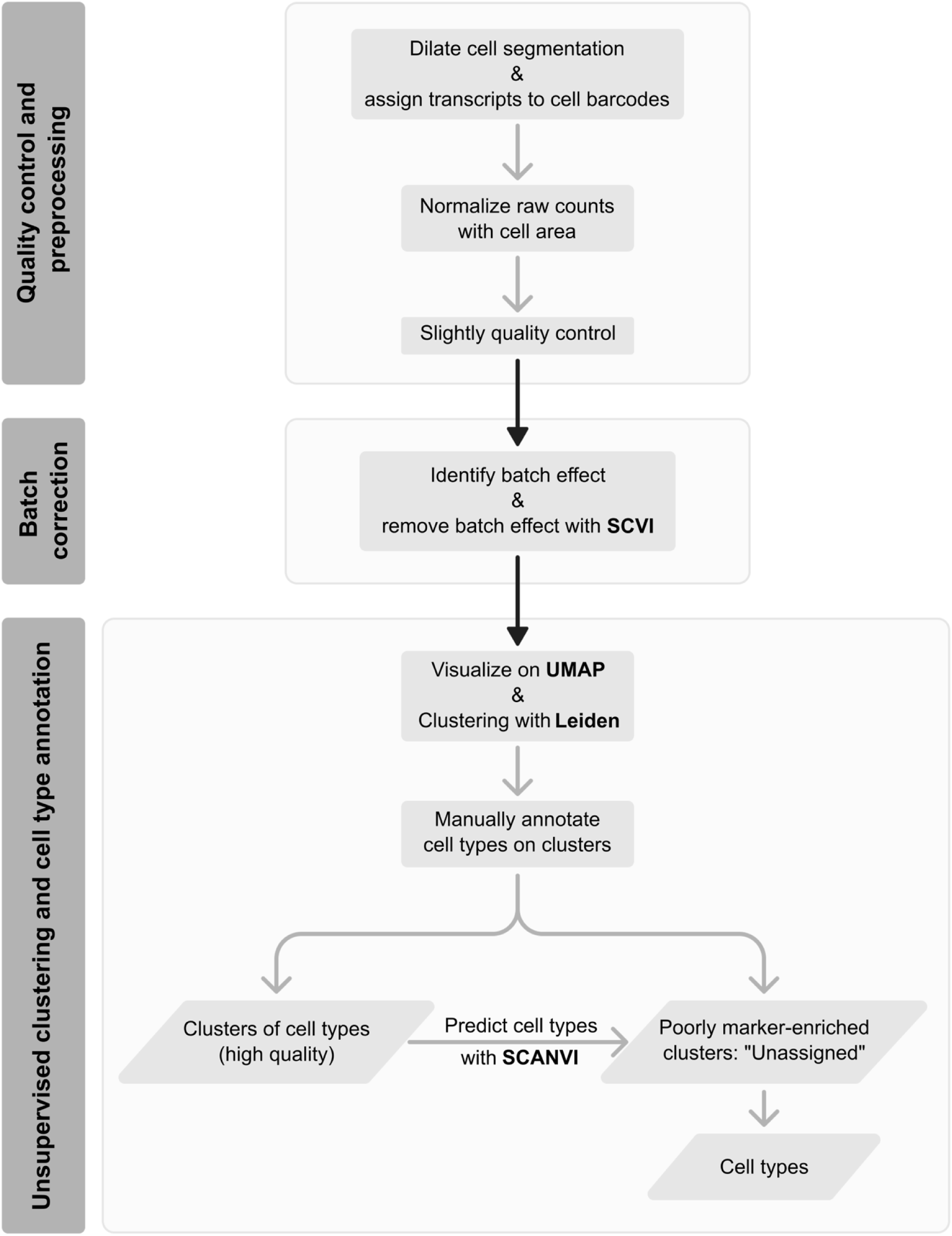
Pipeline for the annotation of cell types using the CosMx dataset. The workflow includes quality control and preprocessing, batch correction and unsupervised clustering and cell type annotation applied for the study.

### Mapping transcripts to cells

Transcripts were mapped to cells to extract their gene expression. Using the cell segmentation provided by NanoString, we expanded the boundaries of each cell, dilating them up to 10 pixels or until they met another cell’s boundary. Transcripts that did not belong to any cell were removed from further analysis. This process resulted in an expression matrix of n_cells x n_genes, where the expression values represent the transcript counts.

### Quality control and preprocessing

scRNA data that captures whole transcriptomics conventionally applies filtering parameters to remove doublets (using sophisticated tools or by applying an upper threshold for unique molecular identifier counts or genes per cell) (8,9). By contrast, CosMx data included a pre-designed panel of 1000 or 6000 genes tailored to specific tissues and user requirements. Cells with fewer than 20 transcript counts were excluded to preserve as many properties as possible. Low-quality cells (if detected) were identified with batch correction and cell type annotation in the latter steps.

Transcript counts were normalized by dividing them by the cell areas, calculated in pixels from the dilated segmentation. We then multiplied all the values by the mean area of all cells to bring the counts back to their original magnitude. Finally, we applied natural log transformation to all expression values. These log - normalized values, instead of log - raw transcript counts, were used for all downstream analyses.

### Identifying and correcting batch effects

We applied a default principal component analysis (PCA) and uniform manifold approximation and projection (UMAP) pipelines on the dataset to check the existence of batch effects between samples (10,11). We employed BioTuring Lens software to perform PCA (default: 50 principal components), KNN (K Nearest Neighbors, default: 30 neighbors), and to generate UMAP space.

After recognizing the presence of batch effect, batch removal was performed by excluding negative control genes, identified by the prefix “NEGPR”. Next, to reduce background noise, we decided to remove roughly 20% of the genes. We utilized the Scanpy package (12) with the “*cell_ranger*” flavor and the previously identified batch information as the “*batch_key*” to select the top 900 highly variable genes (HVGs). These parameters help ensure that differences between batches are also accounted for in selecting top HVGs, where only the normalized variance (z-score) of each gene is considered by default.

Next, a neural-network-based method, single-cell variational inference (scVI) (7), was utilized to remove batch effects. Briefly, the transcript counts of each cell were encoded through a nonlinear transformation into a low-dimensional, batch-corrected, latent embedding. The Variational Auto-Encoder (VAE) model was built with the following parameters: *n_latent = 20, gene_likelihood = “nb”, use_layer_norm = “both”, use_batch_norm = “none”, encode_covariates = True, dropout_rate = 0.2, n_layers = 2.* We then trained the VAE model with a maximum of 400 epochs.

### Unsupervised clustering and cell type annotation

The scVI embedding was loaded onto the BioTuring Lens platform for generating UMAP space and performing clustering (*n_neighbor=50*). To simulate the hierarchical levels of cell types, we ran Leiden clustering at resolutions of 0.5, 1.0, 1.5, and 2.0. We then manually mapped clusters to cell types using the well-established canonical cell type markers in the human kidney, including *PECAM1* for endothelial cells, *NPHS1* for glomerular podocytes, *LRP2* for proximal tubules, *IGHG1* for B cells/plasma cells, and *LYZ* for mononuclear phagocytes (MNP) as previously reported by Balzer et al. (2) (complete gene panel is listed in the Table S1). For annotating sub-cell type classes like principal cells or intercalated cells, we first subset into their major class - connecting tubules, and then re-ran UMAP and clustering for annotation.

For the clusters that had low transcript counts, low number of expressed genes, lacked enrichment in any specific markers, or expressed more than one class of marker genes, we labeled those as “unassigned” clusters, categorizing them as “low-quality cells”. They were labelled by employing scANVI (single-cell Annotation using Variational Inference) (13) with reference from high-quality cell type annotation from the same dataset.

## Results

### Normalization by cell area improves the specificity and sensitivity of identifying differentially expressed genes

Conventional normalization methods for scRNA-sequencing gene expression count data involve scaling based on the total detected gene counts within a cell, commonly known as size factor normalization, relative counts normalization, or library size normalization. However, this approach did not hold for targeted gene panels, such as the 1000-plex gene panel in the CosMx dataset. For image-based platforms such as CosMx, the area of a single cell contributes to the detection rate, with more transcripts detected in larger cell areas. Therefore, we normalized our data by dividing the transcript counts of each cell by its area, calculated in pixels.

We evaluated the new normalization method by performing differential expression analysis (DEG) between the glomerulus and other kidney regions, focusing on genes with well-established roles in glomerular function. Glomerular regions were manually annotated based on CosMx structural images. Evaluating gene list includes *NPHS1* (gene encodes nephrin, a podocyte slit diaphragm protein important in maintaining the glomerular filtration barrier) (14), *FGF1* (fibroblast growth factor 1) (15), *VEGFA* (vascular endothelial growth factor A) (16), *EMCN* (gene encodes endomucin) (17), *PDGFB* (platelet-derived growth factor beta) (18) and *PDGFRB* (platelet-derived growth factor receptor beta) (19). The new normalization method yielded higher log2 fold changes (log2FC) and lower p-values for all six genes, suggesting improved specificity compared to standard log transformation of raw counts (Table 1).

**Table 1.** Comparison of differentially expressed genes in glomerular regions compared to other areas.

| Ctrl_1 and DKD_1-3: Glomerulus vs others |  |  |  |  |
| --- | --- | --- | --- | --- |
|  | Log (normalized) |  | Log (raw-count) |  |
|  | p_value | Log2(FoldChange) | p_value | Log2(FoldChange) |
| NPHS1 | 6.32E-172 | 3.338992882 | 1.38E-170 | 3.15E+00 |
| FGF1 | 0 | 3.86813974 | 0 | 3.692408059 |
| VEGFA | 1.44E-90 | 2.501898801 | 2.85E-87 | 2.34E+00 |
| EMCN | 1.32E-160 | 2.718140343 | 5.13E-159 | 2.58E+00 |
| PDGFB | NaN | NaN | NaN | NaN |
| PDGFRB | 9.31E-06 | 0.7426346519 | 1.53E-05 | 5.88E-01 |
| Ctrl_2 and DKD_2-3: Glomerulus vs others |  |  |  |  |
|  | Log (normalized) |  | Log (raw-count) |  |
|  | p_value | Log2(FoldChange) | p_value | Log2(FoldChange) |
| NPHS1 | 0 | 3.583162993 | 1.38E-170 | 3.15E+00 |
| FGF1 | 0 | 3.40931084 | 0 | 3.692408059 |
| VEGFA | 0 | 3.516809585 | 2.85E-87 | 2.34E+00 |
| EMCN | 0 | 3.423994527 | 0 | 3.299949556 |
| PDGFB | 0.03594627908 | 0.3939787662 | 0.04139500941 | 0.2543612656 |
| PDGFRB | 0.01041549212 | 0.3079504968 | 0.01562241408 | 0.1912055279 |

Additionally, we assessed the entire differentially expressed gene (DEG) output under two normalization strategies using z-scores, p-values, and log2FC values. Across all metrics, the new normalization method demonstrated stronger statistical signals, with higher absolute scores of log2FC values compared to raw count normalization (Tables S2, S3). Moreover, when applying a p-value threshold of <0.05 and log2FC ≥ 1, the new method identified a greater number of differentially expressed genes than log - raw counts (Figure S2), further supporting its enhanced sensitivity.

### Single-cell variational inference (scVI) removal of batch effects improves cell clustering and annotation

Batch effects are an inevitable challenge when integrating datasets from different sources. To address batch variations caused by sample differences, we divided our four samples into two batches containing two control samples and two diabetic patient biopsies classified as class III DKD based on the Renal Pathology Society classification (Figure 2a) (6). We confirmed the presence of batch effects by applying a standard scRNA analysis pipeline and observing how cells from different samples clustered. As expected, cells from the four samples occupied distinct regions in the uniform manifold approximation and projection (UMAP) space (Figure 2a), confirming the need to address batch effects in subsequent analyses. We then employed scVI to remove the batch effects from four samples. Cell types were labelled based on canonical marker genes and the Leiden clustering algorithm on the scVI latent space.

**Figure 2.**
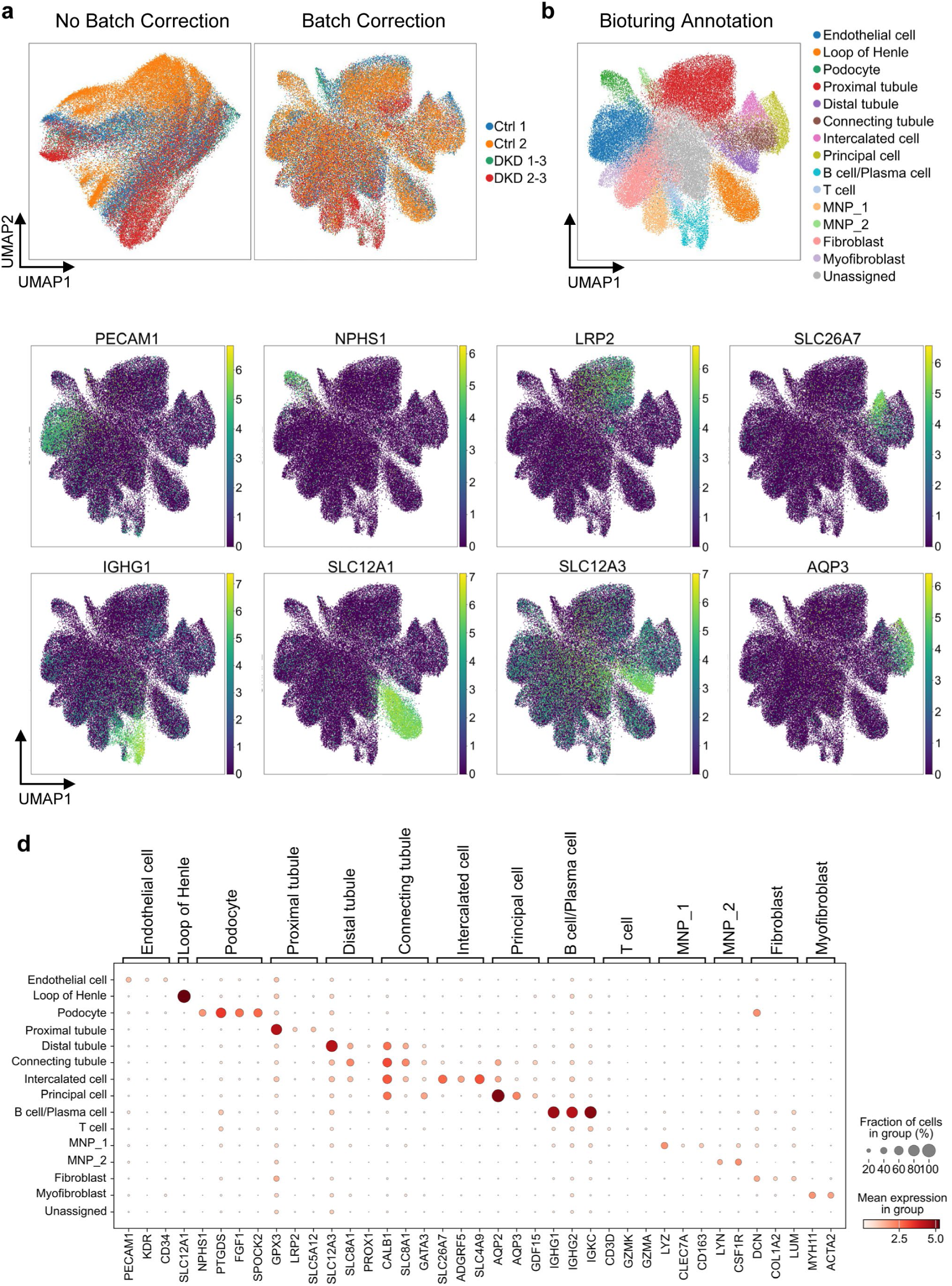
Kidney cell type annotation from single cell spatial transcriptomics. (a) UMAP (Uniform Manifold Approximation and Projection) plots showing 15 cell types in the kidney using the combined spatial dataset for two control (Ctrl 1 and Ctrl 2 with two biopsy proven patients with class III DKD, DKD 1-3 and DK 2-3) before and after batch correction. (b) UMAP plots of the cell clusters for genes encoding known proteins of the kidney following batch correction. (c) UMAP plots of representative genes encoding resident kidney proteins of the individual clusters. (d) Dot plot showing the expression of representative marker genes across the different cell types.

To assess the effectiveness of scVI-learned embeddings, we examined marker gene expression patterns on UMAP. Our annotated cell types, along with their corresponding marker gene expression, are visualized on the same UMAP derived from the scVI latent space (Figure 2b, c). The distinct clustering of marker genes within specific UMAP regions highlights the effectiveness of our batch correction and dimensional reduction process. Dot plots confirmed that the expression of selected marker genes corresponds with cell annotation (Figure 2d). This supports the reliability of clustering-based annotation, particularly in single-cell spatial technologies, where background noise in gene expression is an inevitable challenge.

### Cell-type annotation in kidney tissue resolves structure components of the kidney

In kidney tissue, the most abundant cell types are tubule cells, particularly the proximal tubule cells (20,21). This prevalence often leads to the mislabeling of other cell types due to its dominant marker gene expression over the others. To further evaluate our pipeline, we investigated the glomerulus regions containing endothelial cells, podocytes, and mesangial cells. Following the manual annotation of proximal tubules based on pathological tissue images, we assessed the cell type composition across four samples (Figure 3a, b). Our approach again demonstrated high accuracy, with over 80% of cells in the glomerulus regions annotated as endothelial cells and podocytes (Figure 3a). Additionally, approximately 5% were identified as proximal tubule cells.

**Figure 3.**
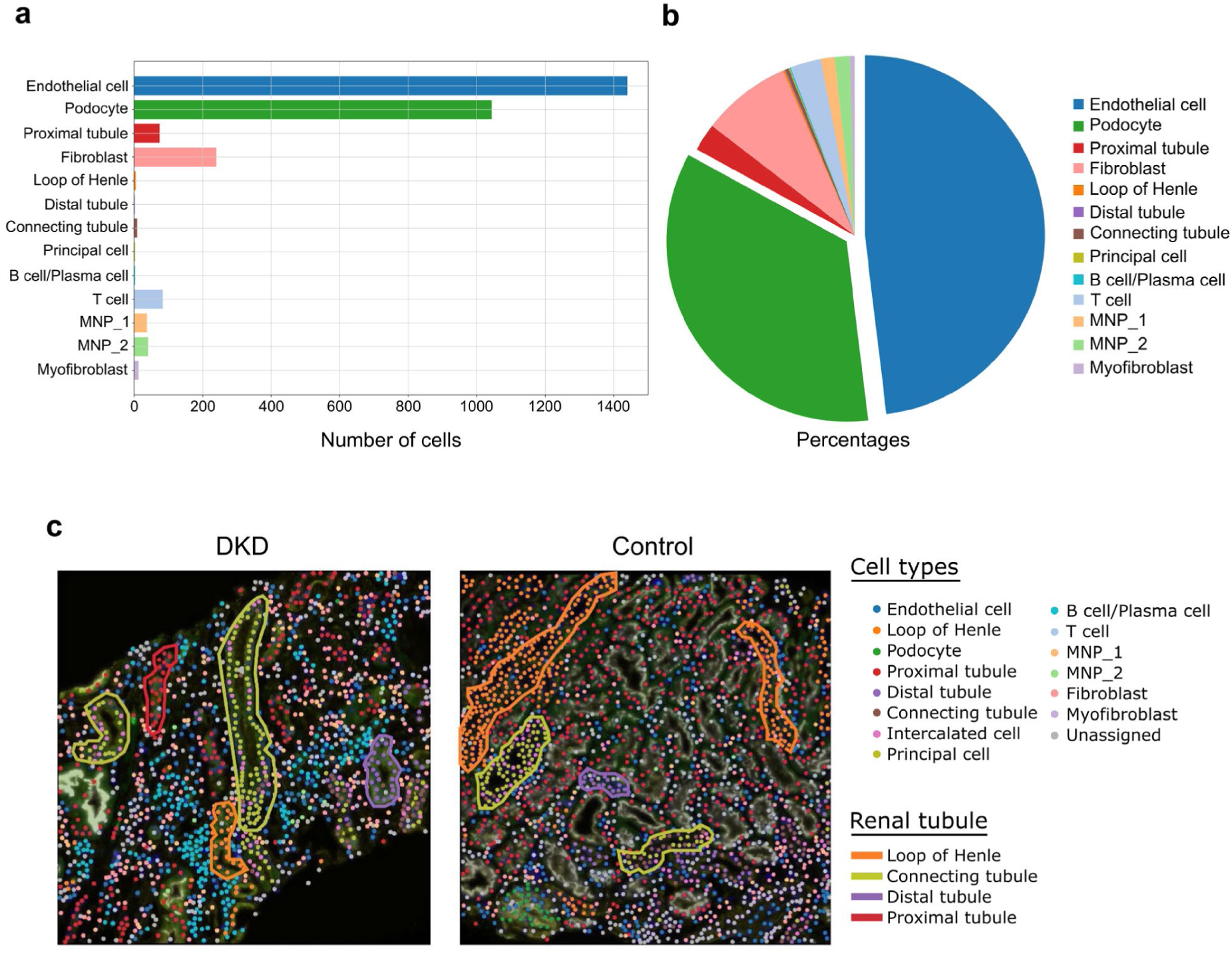
Comparison of cell type annotation in control and DKD tissue. Composition within glomeruli measuring in number of cells (a) and percentage of cell types (b). (c) Comparison of cell types and tubular structures in control and DKD tissue. The cell types were visualized on a pathology image of sample DKD_2-3 and Control_2. “Unassigned” labels are low-quality cells / cells expressing multiple classes of marker genes.

Beyond identifying major cell types of glomeruli, our pipeline also effectively annotated sub-cell types of nephron epithelial cells (Figure 3c). These subtypes were well-separated in the UMAP space and easily annotated using specific marker genes such as *SLC12A3* and *SLC8A1* (distal tubule), *SLC5A12* and *LRP2* (proximal tubule), *SLC12A1* and *S100A2* (Loop of Henle), *AQP2* and *AQP3* (principal cell), and *SLC26A7* and *ADGRF5* (intercalated cell) (Figure 2b). Visualizing cell type annotation on the pathological slides, we observed distinct spatial locations for the loop of Henle (orange), distal tubule cells (purple), and proximal tubule cells (red). The connecting tubules (yellow-green), consisting of principal cells and intercalated cells, were also co-located in specific regions (Figure 3c).

### Analysis of cytokine expression in immune cells and resident kidney cells

After optimizing our cell-type annotation, the differences between DKD and control samples were compared by visualizing cell type composition. We observed an increase in immune cell populations in DKD, particularly B cells (Figure 4a). As expected, markers of B cells were low in control kidneys (Figure 4a) (22). Infiltration of immune cells was found in the glomeruli and interstitium of DKD kidney biopsies, as previously reported (23). Differential expression analysis was conducted between DKD and control samples to select for the top 10 differentially expressed genes (Figure 4b). These genes included *TYK2*, which encodes for a Janus kinase responsible for cytokine-mediated intracellular signaling (24), as well as *AZU1*, encoding azurocidin 1/heparin-binding protein which is known to have functionality as an inflammatory mediator in sepsis (25), and *DUSP5*, which also has a role in hypertension related kidney injury (26). *TYK2* showed increased expression in mononuclear phagocytes population 1 (MNP-1) and T cell immune populations, as well as increased expression in proximal tubule renal populations (Figure 4b). *DUSP5* and *AZU1* also showed increased expression in MNP-1, MNP-2, T cell, and proximal tubule cell populations (Figure 4b,c). *GPX3* (encodes glutathione peroxidase 3, a selenoprotein enzyme secreted by the PCT that acts as an antioxidant) and mitochondrial-associated genes such as *MTRNR2L1* had decreases in expression across cell populations (Figure 4b,c). Taken together, these results validate the increase of immune cell populations in DKD with increased expression levels of genes related to immune signaling and inflammation pathways.

**Figure 4.**
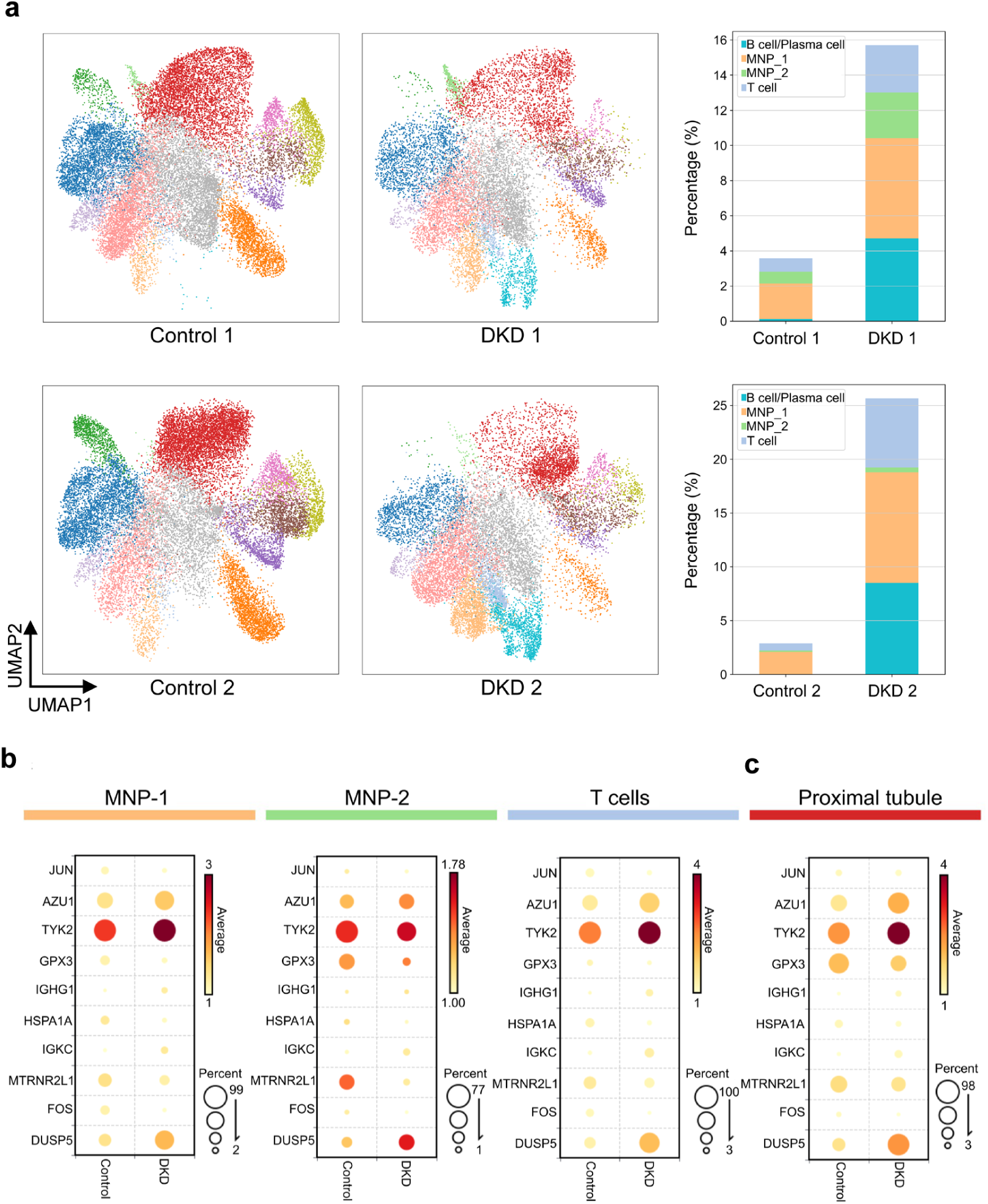
Enrichment of immune cells in DKD. (a) UMAP plots and proportion of immune cell populations in control and DKD tissue. Expression of top 10 differentially expressed genes in healthy controls vs DKD compared in immune (b) and resident cell (c) populations.

### Assessment of injury to podocytes and proximal tubules in diabetic kidney disease

A hallmark indicator of DKD is injury to podocytes and proximal tubular cells. By UMAP comparison, we observed a reduction in the podocyte population in DKD (Figure 5a, b). Subsequently, the expression of *NPHS1* as a marker gene for podocytes was seen to be reduced in the podocyte populations in DKD (Figure 5c). With this data suggesting that podocyte populations were reduced in DKD, the impacts on proximal tubule populations were investigated next. The expression of the marker *HAVCR1* identified injured proximal tubule populations. By comparing the ratio of injured proximal tubules to healthy proximal tubules, an increase in injured proximal tubules can be seen in the diabetic tissue compared to the control (Figure 5d, e). The expression of *HAVCR1* was increased in DKD compared to control proximal tubules, supporting increased tubule injury (Figure 5f). Spatial localization of *VCAM1* and *HAVCR1* for injured proximal tubules, *LRP2* for all proximal tubules, and *NPHS1* for podocytes was compared with cell typing (Figure 5g). Increased expression of *HAVCR1* and *VCAM1* and co-localization with cells labelled as iPTs was observed in DKD tissue (Figure 5g). An increase in the infiltration of B cells and T cells was also observed in DKD compared to control (Figure 5g). In addition, a reduction of *NPHS1* expression was also seen, with the glomerulus structure visible in the DKD tissue showing a reduction in podocytes, indicating glomerular damage (Figure 5g). Taken together, these results support the identification of characteristic podocyte and tubular injury observed in DKD.

**Figure 5.**
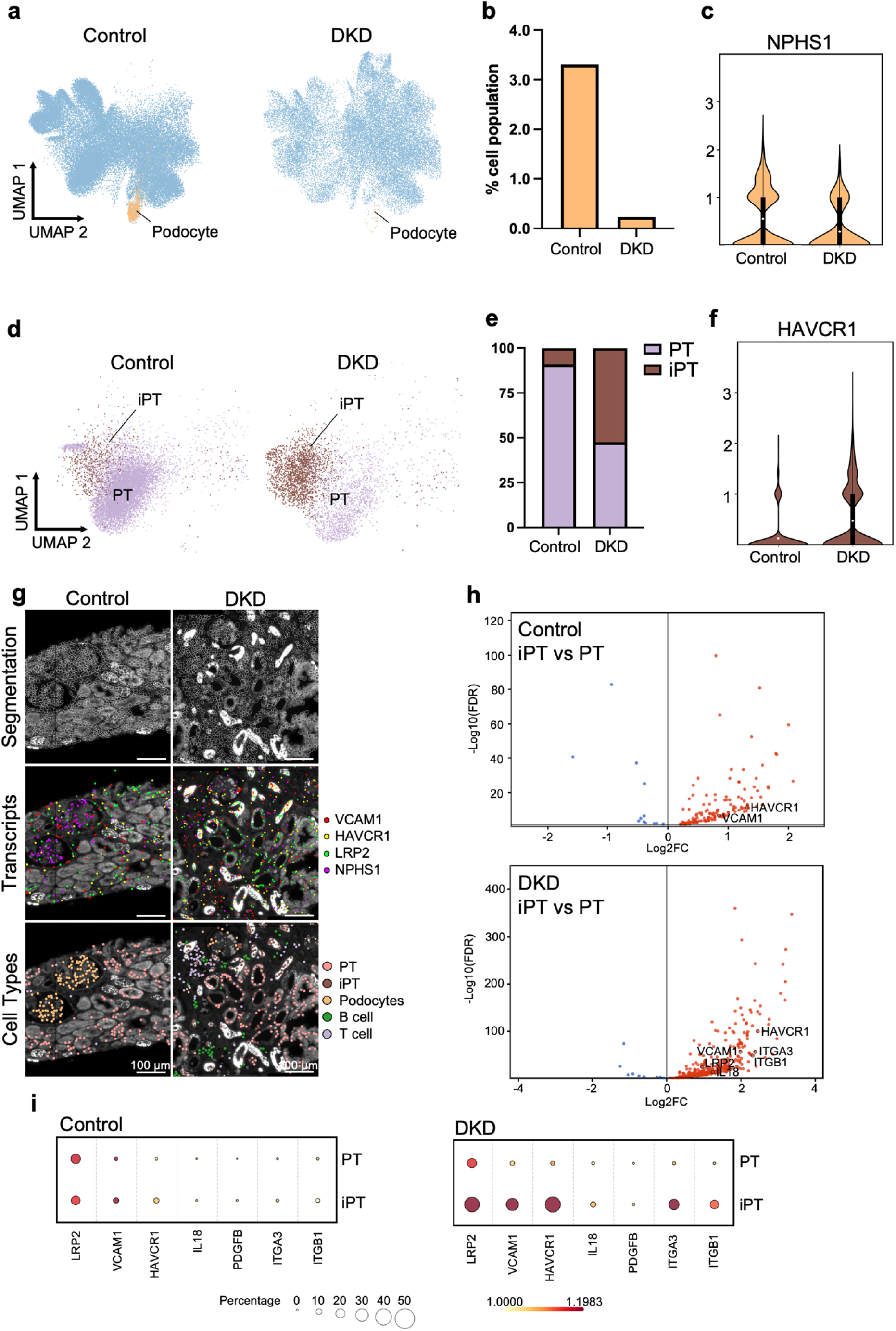
Distribution of podocyte and proximal tubule populations in DKD. (a) UMAP plots comparing proportion of injured podocytes labelled with *NPHS1* between control and DKD tissue. (b) Reduction of podocytes in DKD compared to control tissue. (c) Violin plot of NPHS1 expression (podocyte slit diaphragm gene) between control and DKD podocyte populations. (d) UMAP plots comparing proportion of injured proximal tubules (iPT), labelled with *HAVCR1*, to healthy proximal tubules (PT) between control and DKD tissue. (e) Composition ratio of injured proximal tubules to healthy proximal tubules in control and diseased tissue. (f) Violin plots comparing differential expression of *HAVCR1* proximal tubule injury gene between control and DKD tissue. (g) Tissue visualization of PanCK staining showing cell segmentation, marker gene expression, and tissue resident cell labelling. (h) Expression changes of hallmark diabetes and proximal tubule injury markers. (i) Dot plots of gene expression for the indicated genes control and DKD tissue in PT and iPT.

Changes in hallmark diabetes-associated genes were analyzed in the proximal tubules to further validate our analysis pipeline’s disease modelling capability (Figure 5h). In the control tissue, increased expression of *HAVCR1* and *VCAM1* was observed in the iPT population compared to PTs (Figure 5g). The same held true when comparing the iPT and PT populations in the DKD group, but other notable genes, such as *IL18*, *ITGA3*, and *ITGB1*, were also found to be upregulated (Figure 5h, i). There was no change in the genes *IL18*, *ITGA3*, and *ITGB1* in the control group, indicating the changes in these genes were associated explicitly with DKD (Figure 5i). These results are reinforced by similar trends being observed in the data set analyzed by Abedini et al. (1). Our pipeline presents an accurate cell type annotation for kidney-tissue spatial transcriptomics data, enabling confident analysis of changes in disease states for kidney diseases, not limited to DKD.

## Discussion

In this study, we developed and validated a robust workflow for accurate cell type annotation in single-cell spatial datasets, particularly focusing on kidney tissue from CosMx technology. Improved precision in cell type identification helps to enable critical insights into kidney function, disease mechanisms, and cellular composition across different conditions. The integration of our pipeline into the platform BioTuring SpatialX (Lens V2.0) will provide researchers with limited coding or bioinformatics skills to study changes in kidney diseases.

Our approach has demonstrated significant improvements in identifying key genes while effectively filtering out noise resulting from technical artifacts, such as inaccuracies in cell segmentation. This method resulted in the identification of more significant genes compared to using raw-count data, showcasing the utility of our pipeline in producing cleaner, more biologically relevant results. Although platforms like CosMx provide control probes such as ERCC and NEGPR, which could be used for normalization, we have not yet leveraged them. Incorporating these control probes would further improve data normalization, where positive control probes could serve to normalize gene expression across cells and negative control probes to remove background expression noise. Such advancements would likely lead to more accurate cell type annotations and yield more robust and validated analytical outcomes, improving the precision of downstream analyses like cell type identification and differential expression studies.

The typical pipeline for batch correction developed for scRNA-seq practice involves data normalization, dimensionality reduction by PCA, and application of Harmony for batch effect removal (27). The limitation of this approach is that the PCA’s performance relies heavily on the assumption of linear relationships between genes, which is achieved through the normalization step. However, unlike scRNA-seq data, single-cell spatial data such as CosMx can be impacted by background noise in gene expression (28). This false positive issue, often due to gene expression from neighboring cells or extracellular RNA, complicates the isolation of the target cell’s signal in tissue sections. Since Harmony operates in a lower-dimensional space, it may struggle to retain sufficient biological signals in highly sparse or noisy datasets. Furthermore, because there is currently no standardized normalization method for CosMx technology, the scRNA-seq pipeline is not directly applicable to this spatial technology.

We show that scVI, which works directly with raw counts, offers a more reasonable approach for single-cell spatial data. scVI leverages variational autoencoders (VAEs) to model the underlying probability distribution of count-based data. This allows scVI to effectively separate biological signals from background noise, addressing both the sparsity and false-positive issues seen in CosMx technology. scVI then learns a latent representation and performs batch effect removal based on the true biological variability, making it more effective for single-cell spatial technology.

To analyze the effectiveness of our annotation in identifying disease-induced tissue changes, healthy control tissue was compared to diabetic kidney disease biopsies. While the mechanisms driving DKD progression are still largely unresolved, our study highlights the regional changes in podocyte and tubular injury markers, as well as regional immune cell recruitment and inflammation. Here, we demonstrated the ability to accurately annotate kidney cell types, allowing for disease modelling that agreed with these previously published findings, highlighting immune cell recruitment and proximal tubule and podocyte injury (29). Additional single-cell analysis has uncovered hallmark upregulated genes observed in DKD progression(1). Our dataset showed similar expression trends in hallmark DKD markers such as *IL18*, *ITGA3* and *ITGB1*, as demonstrated previously. Our spatial transcriptomics pipeline presented here demonstrated the ability to identify trends in diabetic kidney disease tissue similar to those previously outlined in the literature. However, this data set was limited by sample size, only including two control and two disease tissue samples. In the future, expanding the sample set could enable novel insights into molecules of interest related to DKD progression. Additionally, with the ever-evolving landscape of spatial transcriptomic platforms, improvements in the number of genes that can be sampled in one experiment will improve cell type annotation and offer deeper insight into possible mechanisms of disease progression.

Despite the robustness of our pipeline, several limitations should be noted. Although we excluded low-quality cells by filtering outliers based on the distribution of detected genes and transcript counts, the filtering parameters were not rigorously defined, potentially affecting the reproducibility and accuracy of the results. In detail, after the whole pipeline, there are still clusters where cell types could not be confidently identified due to a lack of marker gene enrichment and we also labeled them as “low-quality”. To temporarily address this, we used scANVI (13) to transfer the labels from our previous annotation to these areas. However, this inevitably reduces the confidence in drawing firm biological conclusions, and we recommend focusing on high-quality clusters for more reliable findings. Furthermore, the current pipeline does not incorporate spatial information, such as cellular coordinates, into the cell type annotation process. Integrating spatial data could enhance the accuracy of annotations and offer deeper insights into tissue architecture and cellular interactions, thereby improving biological interpretations.

In summary, despite some remaining limitations, our pipeline proves accurate and effective in analyzing CosMx data and utilizing this annotation for disease research. Given that many other single-cell spatial technologies, such as MERSCOPE (Vizgen) and Xenium *in situ* (10X Genomics), face similar challenges with limited gene panels and background noise expression, our pipeline holds promise for broader application across these platforms and different tissues. With the inevitable introduction of the larger whole-transcriptome spatial analysis panels, our pipeline acts as a launching point for more robust analyses that will soon become available.

## Acknowledgements

This research was supported by the NanoString CosMx SMI Technology Access Program with the team of Yan Liang, Yi Cui, Emily Killingbeck, Evelyn Metzger and Andy Nam, who provided access to the spatial molecular imager and raw data. Support for infrastructure and technical was provided by the Live Cell Imaging Facility and the Biobank for the Molecular Classification of Kidney Disease at the Snyder Institute for Chronic Diseases, University of Calgary.

## Author Contributions

SV created the pipelines, analyzed data, wrote the manuscript and prepared figures. KM prepared figures, analyzed data and wrote the manuscript. HD provided bioinformatics support. KC and GA organized the human data and processing of tissue. DAM provided reagents, access to human tissue and editing of the manuscript. TP supervised the bioinformatics and integration of the workflow for BioTuring. JC conceived of the study, contributed to the writing of the manuscript, analyzed data and supervision of the study.

## Data availability

CosMx datasets generated in this study will be made available for reanalysis upon reasonable request. CosMx datasets will be submitted to Gene Expression Omnibus (GEO, https://www.ncbi.nlm.nih.gov/geo/) before publication.

## Supplementary Tables

Table S1: Kidney marker genes.

Table S2: Traditional Wilcox Rank Sum test. Group 1: glomerulus regions, Group 2: others.

Table S3: Scanpy rank genes group. Group 1: glomerulus regions, Group 2: others.

